# Determinants of Adenine-mutagenesis in Diversity-Generating Retroelements

**DOI:** 10.1101/2020.04.29.068544

**Authors:** Sumit Handa, Andres Reyna, Timothy Wiryaman, Partho Ghosh

## Abstract

Diversity-generating retroelements (DGRs) vary protein sequences to the greatest extent known in the natural world. These elements are encoded by constituents of the human microbiome and the microbial ‘dark matter’. Variation occurs through adenine-mutagenesis, in which genetic information in RNA is reverse transcribed faithfully to cDNA for all template bases but adenine. We investigated the determinants of adenine-mutagenesis in the prototypical *Bordetella* bacteriophage DGR through an *in vitro* system composed of the reverse transcriptase bRT, Avd protein, and a specific RNA. We found that the catalytic efficiency for correct incorporation during reverse transcription by the bRT-Avd complex was strikingly low for all template bases, with the lowest occurring for adenine. Misincorporation across a template adenine was only somewhat lower in efficiency than correct incorporation. We found that the C6, but not the N1 or C2, purine substituent was a key determinant of adenine-mutagenesis. bRT-Avd was insensitive to the C6 amine of adenine but recognized the C6 carbonyl of guanine. We also identified two bRT amino acids predicted to nonspecifically contact incoming dNTPs, R74 and I181, as promoters of adenine-mutagenesis. Our results suggest that the overall low catalytic efficiency of bRT-Avd is intimately tied to its ability to carry out adenine-mutagenesis.

## INTRODUCTION

Adaptation by organisms to novel selective pressures requires variation. While this usually occurs over multiple generations and lengthy time scales, there are two examples of instantaneous adaptation that take place within a single generation. These are the variation of antigen receptors by the vertebrate adaptive immune system and of select proteins belonging to diversity-generating retroelements (DGRs) (1). DGRs are prevalent in the human virome and microbiome, and the microbial ‘dark matter’, which appears to constitute a major fraction of microbial life (2–7). The level of DGR variation greatly exceeds the 10^14-16^ variation of the vertebrate immune system (8). A DGR variable protein with 10^20^ possible sequences has been structurally characterized (9), and one with 10^30^ possible sequences has been identified (10). In the adaptive immune system, variation enables the recognition of novel targets and consequent adaptation to dynamic environments. A similar benefit appears to be provided by DGRs, as documented for the prototypical DGR of *Bordetella* bacteriophage (1). This DGR encodes the bacteriophage’s receptor-binding protein Mtd. Variation in Mtd enables the bacteriophage to adapt to the loss of potential surface receptors by its host *Bordetella*, which happens because of environmental changes or immune pressure (11,12).

DGRs vary protein sequences through a mechanism that is fundamentally different from that of the adaptive immune system and indeed any biological system. Sequence variation by DGRs arises from adenine-mutagenesis, in which genetic information is transmitted faithfully for all bases but adenine (Fig. 1). This occurs during reverse transcription of an RNA transcript that contains the intergenic template region (*TR*), which is nearly identical in sequence to a variable region (*VR*) within a DGR variable protein gene. Misincorporation at adenines during reverse transcription of *TR*-RNA results in adenine-mutagenized *TR*-cDNA, which homes to and replaces *VR*, giving rise to a new protein variant. As shown for the *Bordetella* bacteriophage DGR, misincorporation at *TR-*RNA adenines occurs at an astonishingly high frequency of 50% (1,13,14).

**Figure 1.**
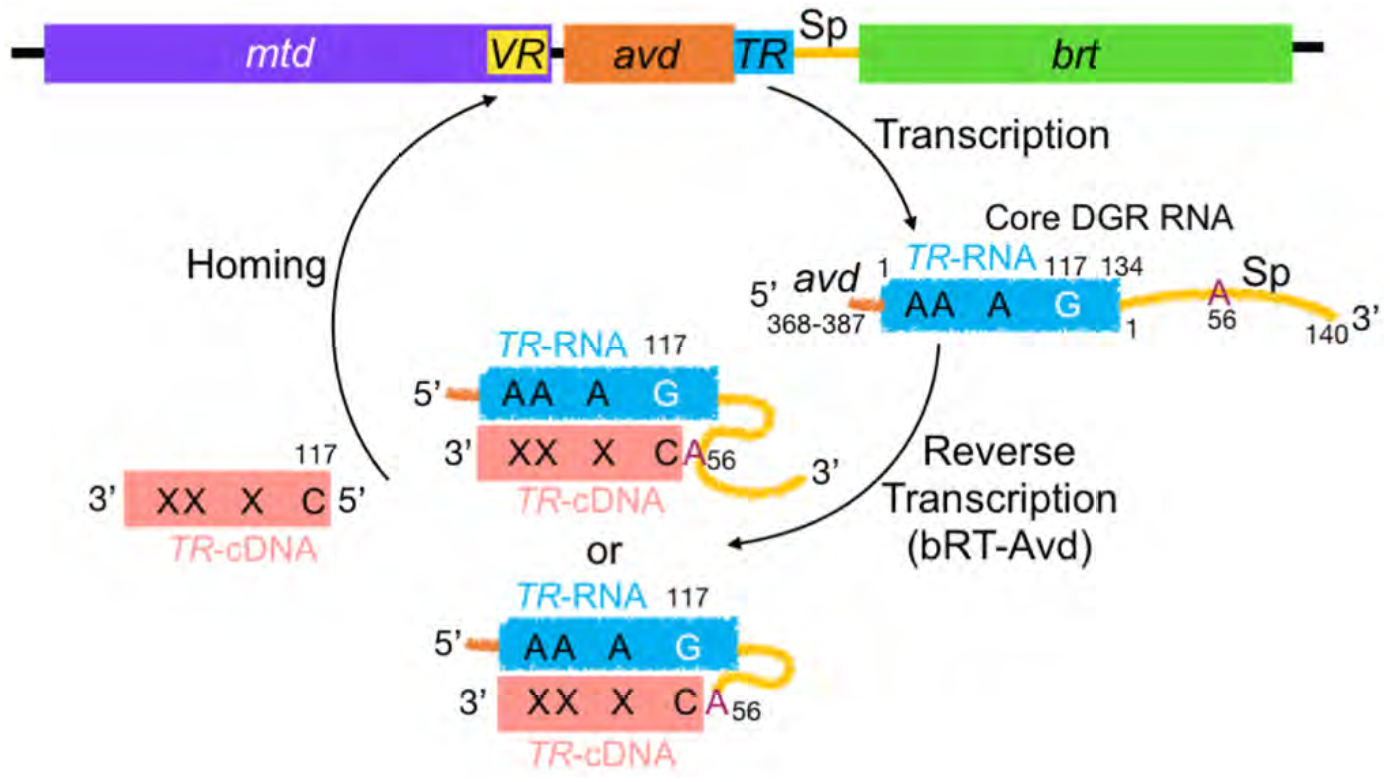
Diversity-generating Retroelement. The *Bordetella* bacteriophage DGR consists of the variable protein gene *mtd*, which contains a variable region (*VR*); *avd* (Accessory Variability Determinant); an intergenic template region (*TR*); an intergenic spacer (Sp) region; and a reverse transcriptase (*brt*). An RNA transcript that contains *TR* and 5′ and 3′ sequences from *avd* and Sp, respectively, is reverse transcribed and adenine-mutagenized by the bRT-Avd complex. Synthesis of *TR-*cDNA is primed by the RNA from Sp A56, either through an internal 2′-OH or a terminal 3′-OH following cleavage of the RNA. *TR* G117 is the first position reverse transcribed, and cDNAs extend to *TR* 22-24 or just into *avd*. *TR*-cDNA homes to and replaces *VR*, involving either the covalently linked RNA-cDNA molecule or the cDNA alone.

We recently reconstituted adenine-mutagenesis *in vitro* (14). This reconstitution showed that a complex formed by the *Bordetella* bacteriophage DGR reverse transcriptase bRT and the associated pentameric Accessory Variability Determinant (Avd) protein (15), along with a specific DGR RNA, are necessary and sufficient for adenine-mutagenesis (Fig. 1). The DGR RNA contains the 134-nucleotide (nt) *TR* flanked by two functional elements, a short 20-nt segment from *avd* at the 5’ end and a longer 140-nt spacer (Sp) region separating *TR* and *brt* at the 3’ end (14). This RNA is identical to the ‘core’ DGR RNA described previously (14). The mechanistic role of the 5’ *avd* sequence is unknown, but the Sp region has been shown to provide an essential binding site for Avd and to supply the site from which reverse transcription is primed, Sp A56 (13,14). A number of pieces of evidence indicate that the 2’-OH of Sp 56 provides the priming nucleophile, resulting in a branched, covalently-linked RNA-cDNA molecule (14) (Fig. 1). An alternative model involving RNA cleavage at Sp 56 to expose its 3’-OH for priming has been proposed as well (13). The first nucleotide reverse transcribed is *TR* G117, and cDNAs typically extend from there to *TR* 22-24 (~90 nt) or just further into *avd* (~120 nt); the shorter cDNA includes all of the adenines in *TR* whose substitution leads to a coding change.

The bRT-Avd complex is also capable of synthesizing cDNAs from non-DGR RNA templates (14). For this, an exogenous primer is required and only short cDNAs (~5-35 nt) are synthesized. These results indicate that template-priming and processive polymerization are both specific properties of the DGR RNA. Evidence suggests that this is because bRT-Avd and the DGR RNA combine to form a structured ribonucleoprotein (RNP) particle that aligns the priming site at Sp 56 with the reverse transcription start site at *TR* 117, and also maintains interaction between bRT-Avd and the RNA that is conducive to processive polymerization (14). Avd is still required for cDNA synthesis from non-DGR RNA templates, indicating that Avd has a role in catalysis that is independent of its role in binding Sp in the DGR RNA. Notably, cDNAs produced from non-DGR RNA templates are adenine-mutagenized, indicating that adenine-mutagenesis is an intrinsic property of the bRT-Avd complex and independent of the RNA template and the mechanism of priming.

To understand the determinants of adenine-mutagenesis, we characterized reverse transcription of the DGR RNA by bRT-Avd *in vitro*. We found that the catalytic efficiency (*k_cat_/K_m_*) of bRT-Avd for correct incorporation was strikingly low across all template bases, as generally observed for low fidelity polymerases (16), with the lowest occurring for adenine. The catalytic efficiency of misincorporation across a template adenine was only somewhat lower than correct incorporation. Using nucleobase analogs, we identified the C6 position of the purine ring as a key determinant of adenine-mutagenesis. An amine at C6, as in adenine, had no effect on misincorporation, neither increasing nor decreasing it. In contrast, a carbonyl at C6, as in guanine, decreased misincorporation. bRT-Avd was able to incorporate dNTPs across an abasic template site, albeit with a significant incidence of deletions and with only a partial preference for adenine as compared to A-rule polymerases (16). We also found that two bRT amino acids, Arg 74 and Ile 181, promoted adenine-mutagenesis. These amino acids are predicted by an *in silico* model to have counterparts in HIV reverse transcriptase (RT) that nonspecifically stabilize incoming dNTPs. These results provide the first detailed characterization of the nucleobase and protein determinants of adenine-mutagenesis in DGRs.

## MATERIAL AND METHODS

### Protein and RNA

**T**he bRT-Avd complex was expressed in *Escherichia coli* and purified, and the core DGR RNA (*avd* 368 – *TR* 140) was produced through *in vitro* transcription with T7 polymerase and gel purified, both as previously described (14). Mutants of bRT were generated using QuickChange mutagenesis (Agilent), and expressed and purified as bRT-Avd complexes. Mutants of the DGR RNA were also generated through QuickChange mutagenesis.

### RNA with nucleobase analogs

RNA oligonucleotides spanning *avd* 368 to *TR* 26 were chemically synthesized (Dharmacon), with adenine, guanine, or base analogs at *TR* 23 and 24 (or just 24), and gel purified. RNA spanning *TR* 27 to Sp 140 was *in vitro* transcribed, gel purified, and treated with alkaline phosphatase (NEB) according to the manufacturer’s directions at 37 °C for 2 h to remove the triphosphate from the 5’ end. The dephosphorylation reaction was quenched at 80 °C for 5 min to inactivate the phosphatase. A phosphate group was added to the 5’ end of the RNA using T4 polynucleotide kinase and ATP, according to the manufacturer’s directions (NEB). The RNA was then purified by phenol:chloroform extraction followed by a G-25 desalting column. The chemically synthesized RNA oligonucleotide (1.1 μM) and the *in vitro* transcribed RNA (2.2 μM) were annealed in the presence of 1.7 μM splint oligodeoxynucleotide P1 (Table S1), 8% DMSO, and 0.2x T4 RNA Ligase 1 (T4Rnl1) buffer in 250 μL. The splint was annealed to the RNA by heating at 95 °C for 3 min and cooling at 0.2 °C/min to 20 °C. A 250 μL solution consisting of 0.8x T4Rnl1 buffer, 2 mM ATP, 4% DMSO, and 540 units T4Rnl1 (NEB) were mixed with the annealing reaction. The resulting mixture was incubated for 8 h at 37 °C. The sample was then extracted with phenol:chloroform and ethanol-glycogen precipitated. The pellet was resuspended in water and gel purified.

### Reverse transcription reactions and next-generation sequencing

Reverse transcription reactions were carried out with wild-type or mutant bRT-Avd at 37 °C for 12 h, and resulting cDNA was purified, both as previously described (14). The region from *TR* 21 to *TR* 98 was PCR amplified using *Pfu* polymerase from purified cDNA using primers P2 and P3, which have partial Illumina adapter sequences at their 5’ ends (Table S1). For amplification of the region from *TR* 114 to *TR* 117, reverse transcription of RNA-cDNA molecules with primer P2 was first carried out, as previously described (14), and the resulting cDNA was PCR amplified with *Pfu* polymerase and primers P2 and P4, which likewise had partial Illumina adapter sequences at their 5’ ends (Table S1). The reverse transcription step was necessary as *TR* 117 is at the 5’ end of the cDNA and attached covalently to the core DGR RNA at Sp 56. The amplified PCR product was subjected to short-read next-generation sequencing (Amplicon EZ, Genewiz). The quality scores for sequencing reactions were Q30, which is equivalent to an error probability of 1 in 1000. Fastq files generated from next-generation sequencing containing paired-end reads were aligned with the *TR* reference sequence using bowtie2, and output files were sorted and indexed using samtools (parameters in Supplementary Data) (17–19). (Mis)incorporation frequencies in the template region were calculated using IGV (integrative genomics viewer) (20).

Reverse transcription reactions with Moloney Murine Leukemia Virus (MMLV) RT (BioBharati Life Sciences Pvt. Ltd) and HIV RT (Worthington Biochemical) were carried out according to the manufacturer’s directions using primer P5 (Table S1). PCR amplification of cDNAs was carried out with *Pfu* polymerase and primers P2 and P3 (Table S1).

### Quantitative PCR

Quantitative PCR (qPCR) was carried out on cDNA reverse transcribed by bRT-Avd from the core DGR RNA (25 μL reaction). The cDNA was purified as previously described (14) and amplified in the presence of 1x SYBR Green 1 dye (Thermo Fisher) using primers P6 and P7 (Table S1). The reaction was performed on a Bio-Rad CFX Connect Real Time System apparatus. A standard curve was generated using chemically synthesized single stranded DNA (*TR* 1 to *TR* 117, HPLC purity of 99.9%), with template concentrations in the range of 100 fg – 1 ng. qPCR reactions were performed in triplicate, and the cycle threshold (Ct, the cycle number at which the fluorescence due to the reaction crosses the fluorescence background threshold) for each reaction was determined using CFX Maestro Software. The average of Ct was plotted against the log of the DNA mass, and a linear fit from the plot was used to calculate the quantity and concentration of cDNA generated in the bRT-Avd reverse transcription reaction.

### Single deoxynucleotide primer extension assay

Oligodeoxynucleotide P^117^ (Table S1) was 5’-[^32^P]-labeled as previously described (14). Wild-type or mutant core DGR RNA (0.5 μM) was mixed with primer P^117^ (0.5 μM), 5’-[^32^P]-labeled P^117^ (0.05 μM), varying concentration of dNTPs, and 20 units RNase inhibitor (NEB) in 75 mM KCl, 3 mM MgCl_2_, 10 mM DTT, 50 mM HEPES, pH 7.5 and 10% glycerol in a 20 μL final volume. The mixture was incubated at 37 °C, and then wild-type or mutant 1 μM bRT-Avd was added. A 2.5 μL aliquot of the reaction was removed at various time points, and quenched by addition to 7.5 μL of an ice-cold solution of proteinase K (1.3 mg/ml) followed by incubation at 50 °C for 20 min. The quenched reactions were incubated with 0.5 μL RNase A/T1 mix (Ambion) at room temperature for 20 min in a final volume of 20 μL. The samples were then ethanol-glycogen precipitated overnight at –20 °C. Samples were centrifuged the next day, and pellets were air dried and resuspended in 20 μL of RNA loading dye. Five μL of the reaction sample was loaded on an 8% denaturing sequencing gel to resolve unreacted and extended primers. The radiolabeled products were visualized by autoradiography using a Typhoon Trio (GE Healthcare Life Sciences), and band densities were quantified using ImageQuant TL 8.1 (GE Healthcare Life Sciences). Background values determined from band densities prior to any reaction were subtracted. The steady-state initial velocity with respect to substrate concentration was fit to the Michaelis-Menten equation using nonlinear regression analysis in GraphPad Prism.

## RESULTS

### Adenine-mutagenesis of *TR*

To characterize adenine-mutagenesis of *TR* by bRT-Avd at fine-scale, we pursued next-generation sequencing (NGS) of cDNAs. The NGS read count of ~100,000 enabled conclusions to be drawn about the distribution of adenine-mutagenesis that were not possible due to the small number of sequences previously available from single clones (~30) (1). The template was the ~300-nt core DGR RNA (14), consisting of the *TR* (134 nt) flanked by upstream *avd* and downstream Sp regions that are functionally essential (Fig. 1) (14). As described above, reverse transcription was template-primed by the core DGR RNA from Sp A56 and initiated from *TR* G117 (14).

To determine the baseline detection level of our methodology, we carried out NGS on cDNA that had not been adenine-mutagenized. For this, we used chemically synthesized, purified single-stranded (ss) DNA that corresponded to *TR* 1-117. During NGS analysis we set the concentration of this ssDNA to be equivalent to that of the cDNA produced by bRT-Avd. The latter was determined through quantitative PCR (Fig. S1a), which showed that 1 x 10^11^ cDNA molecules were synthesized from 6 x 10^12^ DGR RNA molecules. NGS of the non-mutagenized ssDNA yielded an average 0.13% misincorporation frequency, with a range of 0.1-0.7% (Fig. S1b). As a further control, we synthesized *TR-*cDNA from the DGR RNA template using an exogenous primer with the high fidelity reverse transcriptase MMLV RT, which has a reported error rate of ~10^−5^ (21). NGS analysis revealed an average 0.36% misincorporation frequency, with a range of 0.1-2.6% (Fig. S1c). Interestingly, *TR* adenines had the highest average misincorporation frequency (0.8%).

We then carried out NGS analysis on cDNAs synthesized by bRT-Avd. We found through three independent experiments that the average misincorporation frequency across the 22 adenines in *TR* that lead to coding changes was 51.6 ± 2.3% (Figs. 2a and S2a), similar to the level previously observed *in vitro* and *in vivo* (1,13,14). The most frequently misincorporated base across a template adenine was adenine itself (22.2 ± 0.7%), followed by cytosine (17.8 ± 2.8%) and guanine (11.6 ± 0.2%) (Fig. S2a). By comparison, the misincorporation frequency across template uracils, cytosines, and guanines in *TR* was at or near baseline detection levels (0.5 ± 0.2%, 0.3 ± 0.2%, and 1.6 ± 1.2%, respectively) (Fig. S2b).

**Figure 2.**
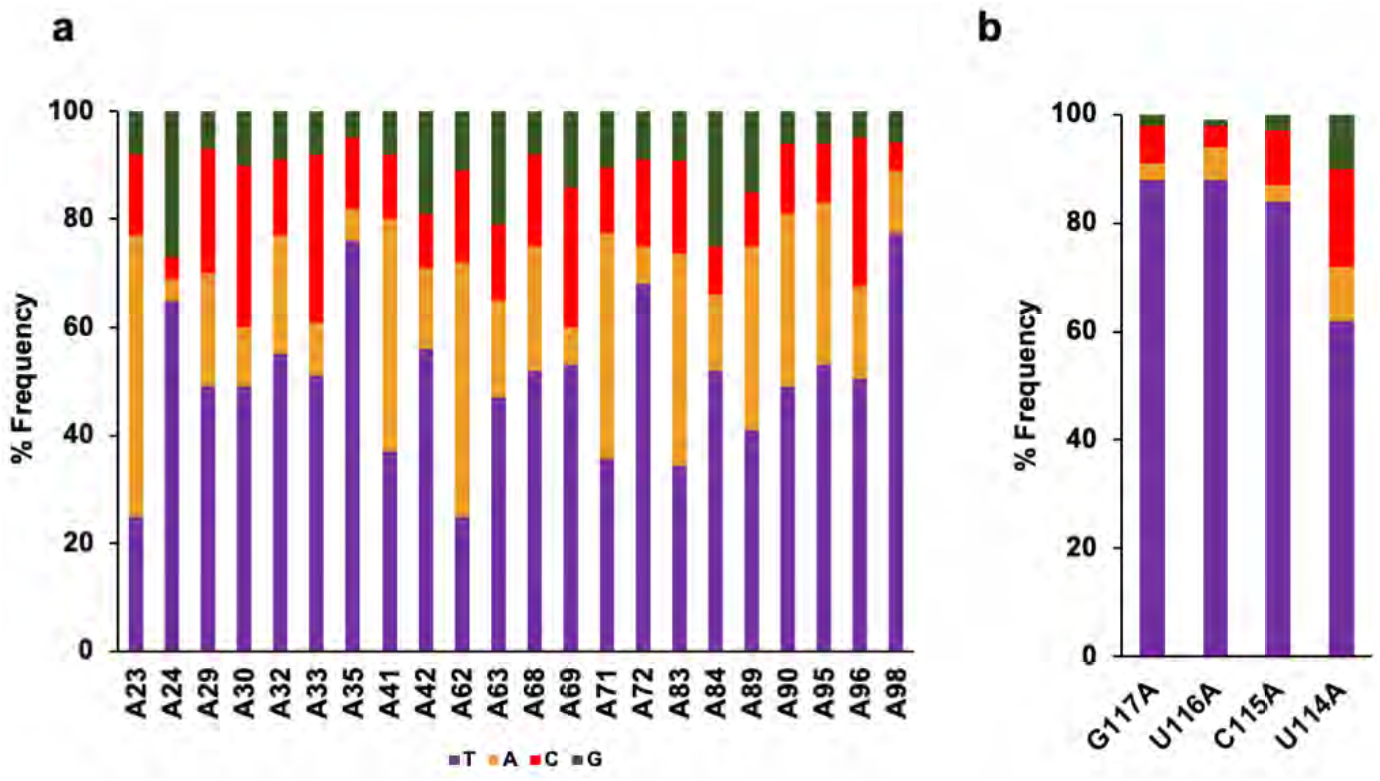
Adenine- mutagenesis. a. Frequency of deoxynucleotides incorporated (thymine, purple) or misincorporated (adenine, orange; cytosine, red; guanine, green) by bRT-Avd across template adenines in *TR*. Results from one of three independent experiments is shown. This color coding is used throughout. b. Frequency of deoxynucleotides (mis)incorporated across template adenines individually substituted at *TR* 114-117.

The misincorporation frequency of bRT-Avd varied widely across individual template adenines (Figs. 2a and S3). *TR* A23 and A62 were especially prone to misincorporation, with frequencies of 72.3 ± 1.9% and 76.3 ± 0.9%, respectively. Notably these adenines are the first bases in AAC codons (for Mtd 344 and 357, respectively), indicating an enhanced potential for amino acid substitution at these positions (22). In contrast, *TR* A35 and A98 were especially resistant to misincorporation, with frequencies of only 28.0 ± 2.8% and 29.9 ± 5.2%, respectively. Equally notably, these adenines are the only ones in their codons (ACG for Mtd 348 and ATC for Mtd 369, respectively), indicating a curtailed potential for amino acid substitution at these codons (22). The differences in misincorporation frequencies were not attributable to any obvious RNA primary sequence patterns.

To determine whether misincorporation also occurred at artificially introduced adenines, we substituted adenines at the initial four positions that are reverse transcribed (*TR* 117-114). These positions are normally occupied by G, C, or U. Misincorporation was evident at all four positions when substituted by adenine. Positions 117, 116, and 115 were somewhat more resistant to misincorporation than the most resistant adenines positions naturally occurring in *TR*, while 114 was within the range observed for naturally occurring adenine positions (Fig. 2b). Thus adenine-mutagenesis can occur outside naturally occurring positions, and similar to naturally occurring positions, the misincorporation frequency is variable.

### Enzymatic Parameters of bRT-Avd

We next examined the rate of single deoxynucleotide addition by bRT-Avd. As misincorporation occurred at a detectable level across *TR* G117A, we used oligodeoxynucleotide P^117^ (Table S1) to prime synthesis from *TR* G117A of the core DGR RNA. We have previously shown that P^117^ primes cDNA synthesis from the natural start site of reverse transcription, *TR* G117, and concurrently inhibits template-primed cDNA synthesis (14). The addition of a single deoxynucleotide to the radiolabeled P^117^ primer was unambiguously detectable (Fig. 3a). In the case of extension with dATP, the template contained both *TR* U116G and G117A substitutions, so as to avoid incorporation of a second dATP across *TR* U116. Using this single deoxynucleotide primer extension assay, we determined the steady-state enzymatic parameters of bRT-Avd for correct incorporation and misincorporation across a template adenine.

**Figure 3.**
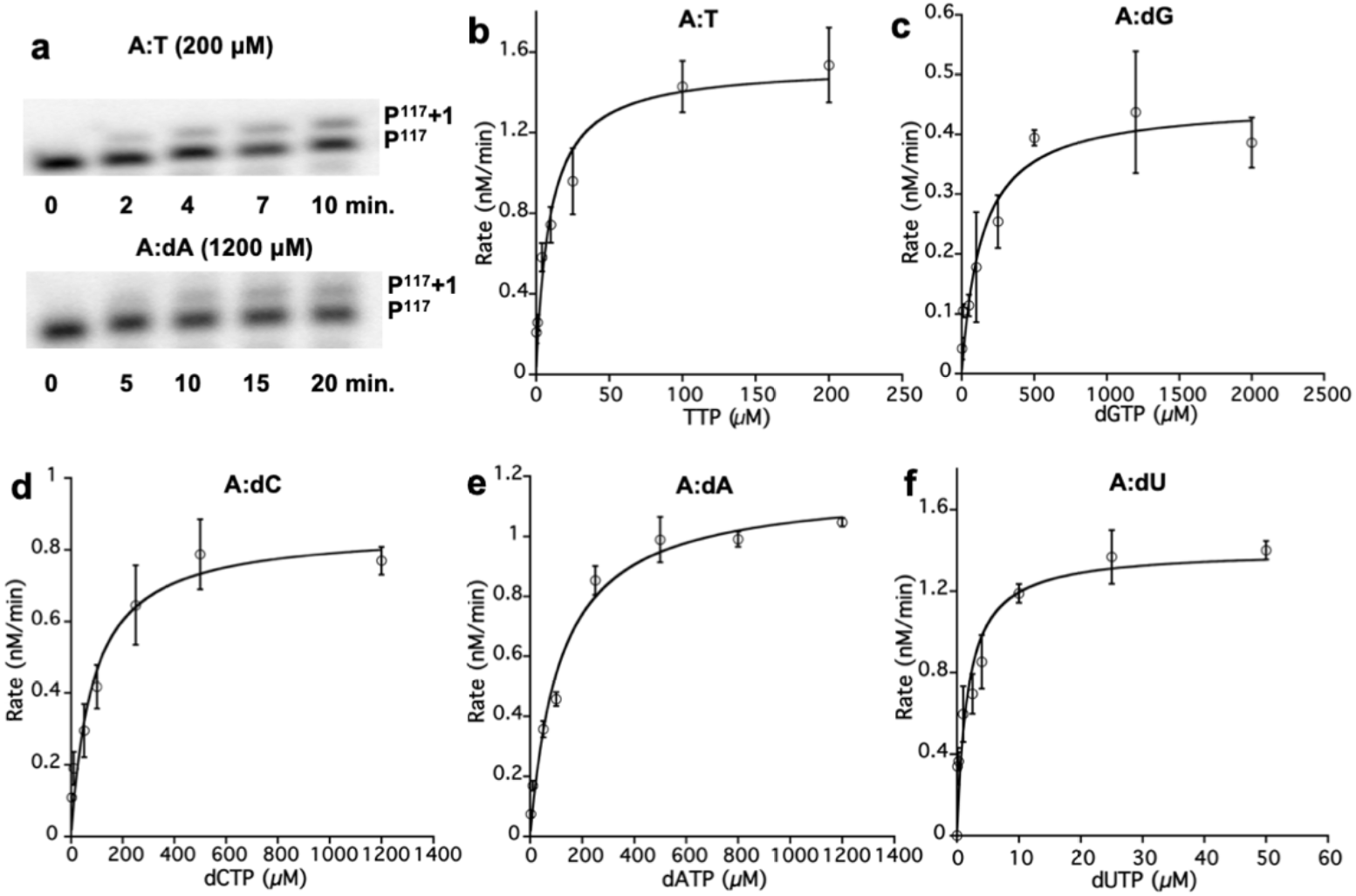
Kinetics of single deoxynucleotide (mis)incorporation. a. Single deoxynucleotide primer extension by bRT-Avd of [^32^P]-labeled P^117^ by TTP (top) or dATP (bottom), as templated by the core DGR RNA containing *TR* G117A. The extended product (P^117^ +1) was resolved from the reactant (P^117^) with an 8% sequencing gel. b-f. Steady-state kinetic characterization of single deoxynucleotide (mis)incorporation of TTP, dGTP, dCTP, dATP and dUTP, across the core DGR RNA containing *TR* G117A (and *TR* U116G, in the case of dATP) by bRT-Avd. The Michaelis-Menten fit is shown, and error bars represent standard deviations from three independent measurements.

This analysis showed that the *k*_cat_ varied little (only up to three-fold) between correct incorporation of TTP and misincorporation of the other dNTPs across the adenine at *TR* 117 (Figs. 3b-f and Table 1). Similarly, the *K_m_* for the incoming dNTP varied over a very small range, with the greatest difference being ~17-fold between dGTP and TTP (Table 1). The most favorable *K_m_* was for dUTP, reflecting the previously noted preference in bRT-Avd for dUTP over TTP (14). The catalytic efficiencies (*k*_cat_/*K_m_*) for the misincorporation of dGTP, dCTP, and dATP were on average 23-fold lower than for the correct incorporation of TTP. The misincorporation ratios (catalytic efficiency for misincorporation to correct incorporation) for dCTP and dATP were 0.05, and 0.02 for dGTP (Table 1). These values corresponded closely to the misincorporation frequencies at *TR* G117A determined by NGS from template-primed reverse transcription (Fig. 2b, 7%, 3%, and 2%, respectively). Thus, template- and P^117^-primed synthesis resulted in similar misincorporation frequencies, which indicated that the enzymatic parameters determined here for oligodeoxynucleotide-primed synthesis were likely to be applicable to template-primed synthesis.

**Table 1.** Steady-state enzymatic parameters for (mis)incorporation by bRT-Avd at *TR* G117A.

| WT | $K_m$ ($\mu\text{M}$) | $k_{cat}$ ($\text{min}^{-1}$) $\times 10^{-3}$ | $k_{cat}/K_m$ ($\mu\text{M}^{-1} \text{min}^{-1}$) $\times 10^{-3}$ | Misincorporation ratio<sup>b</sup> |
| --- | --- | --- | --- | --- |
| A:T<sup>a</sup> | 7.97 ± 2.34 | 11.64 ± 0.71 | 1.46 | - |
| A:dG | 138.03 ± 48.33 | 3.62 ± 0.32 | 0.03 | 0.02 |
| A:dC | 84.35 ± 26.10 | 6.85 ± 0.56 | 0.08 | 0.05 |
| A:dA | 115.25 ± 22.31 | 9.33 ± 0.46 | 0.08 | 0.05 |
| A:dU | 1.73 ± 0.64 | 11.20 ± 0.96 | 6.47 | 4.43 |

<sup>a</sup>Template base:(mis)incorporated base
<sup>b</sup>Misincorporation ratio = $[k_{cat}(\text{incorrect})/K_m(\text{incorrect})] / [k_{cat}(\text{correct})/K_m(\text{correct})]$

We then examined the enzymatic parameters for correct dNTP incorporation across the other template bases at *TR* 117. For template guanine, the wild-type DGR RNA template was used, and for template cytosine and uracil, mutated DGR RNA templates were used (*TR* G117C for the former, and *TR* U116C/G117U for the latter to avoid incorporation of a second dATP). The experiment showed that the *k*cat for correct incorporation varied little among template guanine, cytosine, uracil, and adenine (Tables 1 and 2, Fig. S4). However, there was a major difference in *K_m_*. While the *K_m_* was similar for template guanine, cytosine, and uracil, this value was on average ~36-fold lower than for a template adenine. Similarly, the catalytic efficiencies were nearly identical for these first three template bases but 28-fold greater than for a template adenine.

**Table 2.** Steady-state enzymatic parameters for incorporation by bRT-Avd at *TR* G117, G117C, and G117U.

| WT | $K_m$ ($\mu\text{M}$) | $k_{cat}$ ($\text{min}^{-1}$) $\times 10^{-3}$ | $k_{cat}/K_m$ ($\mu\text{M}^{-1} \text{min}^{-1}$) $\times 10^{-3}$ | Efficiency<sup>c</sup> |
| --- | --- | --- | --- | --- |
| G:dC | 0.26 ± 0.09 | 10.41 ± 0.62 | 40.04 | 27.42 |
| C:dG | 0.17 ± 0.06 | 6.92 ± 0.41 | 40.71 | 27.88 |
| U:dA | 0.24 ± 0.09 | 9.95 ± 0.62 | 41.46 | 28.40 |

<sup>c</sup>Efficiency = $[k_{cat}(\text{correct})/K_m(\text{correct})] / [k_{cat}(\text{wt, A:T})/K_m(\text{wt, A:T})]$

### Nucleobase Determinants of Adenine-Mutagenesis

We next asked which features of adenine promoted misincorporation. To address this, we compared the nearly isosteric features of infidelity-promoting adenine and fidelity-promoting guanine. The differences between A and G primarily occur at the N1, C2, and C6 positions (Fig. 4a). We sought to probe these ring positions using base analogs. For this, short RNA oligonucleotides corresponding to the 5’ portion of the core DGR RNA were chemically synthesized with nucleobase analogs at *TR* 23 and 24, which have misincorporation frequencies of 72.3 ± 1.9% and 39.7 ± 3.3%, respectively. The oligonucleotide was ligated to a longer *in vitro* transcribed RNA corresponding to the rest of the core DGR RNA. To validate this method, we first constructed the ligated core DGR RNA using an oligonucleotide that had adenines rather than analogs at *TR* 23 and 24. The cDNAs produced by bRT-Avd from the ligated core DGR RNA template had a near identical misincorporation frequency as the fully *in vitro* transcribed core DGR RNA template (Figs. 2a and 4b). As further validation, we used the same ligation method to introduce guanines at *TR* 23 and 24, and found almost exclusive incorporation of cytosine by bRT-Avd (Fig. S5), indicating that there is no barrier to correct incorporation at these positions.

**Figure 4.**
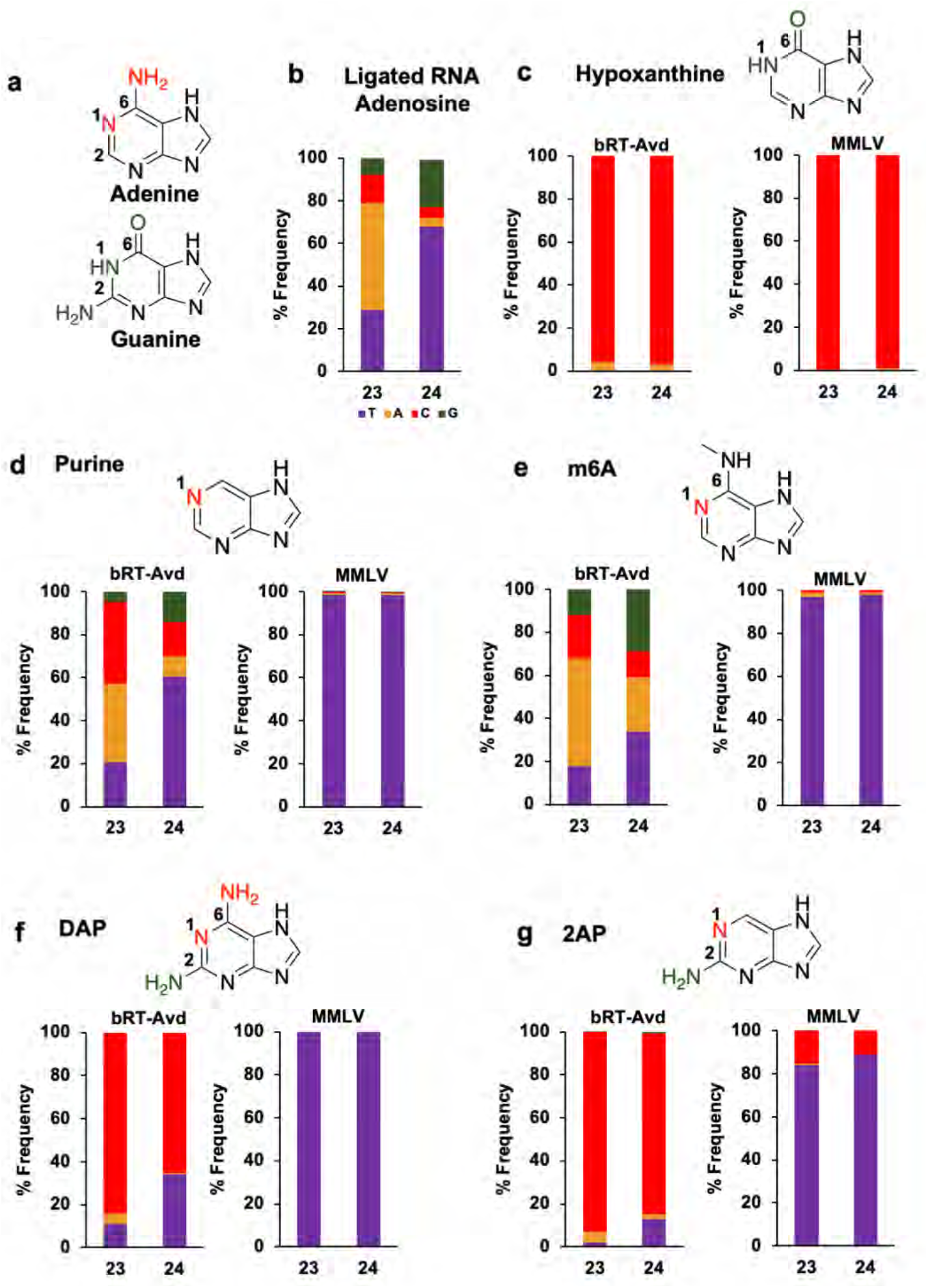
Nucleobase determinants of adenine-mutagenesis. a. Adenine and guanine differ at N1, C2, and C6 positions (red for A, green for G). b. Frequency of deoxynucleotides (mis)incorporated by bRT-Avd at *TR* 23 and *TR* 24 using the core DGR RNA template that had been ligated from chemically synthesized and *in vitro* transcribed sections. The chemically synthesized section contained adenosines at *TR* 23 and 24. c. Top, structure of hypoxanthine. Bottom, (mis)incorporation frequencies of bRT-Avd (left) and MMLV RT (right) with hypoxanthine at *TR* 23 and 24. d. Top, structure of purine. Bottom, (mis)incorporation frequencies of bRT-Avd (left) and MMLV RT (right) with purine at *TR* 23 and 24. e. Top, structure of N6-methyladenine (m6A). Bottom, (mis)incorporation frequencies of bRT-Avd (left) and MMLV RT (right) with m6A at *TR* 23 and 24. f. Top, structure of DAP. Bottom, (mis)incorporation frequencies of bRT-Avd (left) and MMLV RT (right) with DAP at *TR* 23 and 24. g. Top, structure of 2,6-diaminopurine (DAP). Bottom, (mis)incorporation frequencies of bRT-Avd (left) and MMLV RT (right) with DAP at *TR* 23 and 24.

#### N1 and C6

The first nucleobase analog we examined was hypoxanthine, in which the N1 and C6 groups of adenine (N with a lone electron pair and amine, respectively) are substituted with those of guanine (NH and carbonyl, respectively). Hypoxanthine preferentially forms a Watson-Crick base pair with cytosine, and a less stable wobble base pair with adenine (23,24). This preference was verified through reverse transcription with MMLV RT (Fig. 4c). Cytosine was almost exclusively incorporated across template hypoxanthines by MMLV RT. Reverse transcription was then carried out with bRT-Avd. Significantly, bRT-Avd correctly incorporated cytosine with nearly the same frequency, 96-97% (Fig. 4c), as MMLV RT. Adenines constituted the rest. This result showed that an NH at N1 or a carbonyl at C6, or both, decrease misincorporation relative to adenine.

We next asked if the amine at C6 in adenine has an impact on misincorporation. To do so, we used purine (i.e., nebularine), which lacks a substituent at C6, as the base analog. Purine preferentially base pairs with thymine (25), which we verified with MMLV RT (Fig. 4d). With bRT-Avd, template purines led to misincorporation with very similar frequencies as those observed for template adenines (Fig. 4d). This result indicated that the C6 amine of adenine was functionally equivalent to no substituent at this position. Thus, we concluded that an amine at C6 had no impact on misincorporation, neither decreasing nor increasing it.

To probe the C6 position further, we used N6-methyladenine (m^6^A). As previously reported, reverse transcription of m^6^A by MMLV results in the incorporation of thymine (Fig. 4e) (26). However, for bRT-Avd, m^6^A resulted in greater misincorporation, most notably at *TR* 24, where misincorporation increased from 40 to 66% (Fig. 4e). Misincorporation also increased at *TR* 23, although less strikingly, from 72 to 82%. Thus, while an amino group at the C6 position did not impact misincorporation, the bulkier methylamino group at the same position increased misincorporation. Importantly, these results also suggested that the N1 position had little if any role in modulating misincorporation. This is because while adenine and m^6^A are identical at the N1 position, they differed in misincorporation frequency.

Taken together, these results indicated that the C6 but not the N1 purine position was a major determinant of adenine-mutagenesis, and that the C6 carbonyl of guanine decreased misincorporation while the C6 amine was equivalent to having no substituent at this position.

#### C2

The C2 position was probed using 2,6-diaminopurine (DAP), which is identical to adenine except containing an amine at C2 as well (as does guanine). DAP preferentially base pairs with thymine but can also form a wobble base pair with cytosine (27,28). In the case of MMLV RT, exclusive incorporation of thymine was observed (Fig. 4f). However for bRT-Avd, the major species incorporated was cytosine, followed to a lesser extent by thymine and adenine (Fig. 4f). This was the case even though the Watson-Crick base pair between DAP and thymine involves three hydrogen bonds, while the wobble base pair between DAP and cytosine involves only one.

A similar trend was seen for 2-amino purine (2AP), which is identical to DAP but lacks an amine at C6. 2AP forms a base pair with thymine but can also form a wobble or protonated base pair with cytosine (28–30). Both forms involve two hydrogen bonds. MMLV RT preferentially incorporated thymine but also a substantial level of cytosine. In the case of bRT-Avd, incorporation of cytosine was greatly preferred, followed by thymine and adenine (Fig. 4g), similar to the results with DAP (Fig. 4f). The similarity of misincorporation frequencies between DAP and 2-AP (which differ only in an amine at C6) confirmed that an amine at C6 was equivalent to having no substituent at this position.

These results showed that an amine at C2 led to a more homogeneous distribution of incorporated deoxynucleotide, but with those forming a wobble base pair being favored over those forming a Watson-Crick base pair. The exception to this was when the C6 position was occupied by a carbonyl, that is when the template base was a guanine, in which case a Watson-Crick base pair was favored. Thus, these results further emphasized the importance of the C6 position to adenine-mutagenesis.

### Abasic Site and the A-rule

We noticed that adenine was the most frequently misincorporated base across the 22 sites in *TR* (Figs. 2a and S2a). A number of nucleotide polymerases have the tendency to insert an adenine across an abasic template site, which is called the A-rule (31). We investigated whether bRT-Avd follows the A-rule by constructing a core DGR RNA template with abasic sites at *TR* 23 and 24. HIV RT has been documented to tolerate abasic sites (32), and thus we used HIV RT as a positive control. However, tandem abasic sites led to a high incidence of deletions at these and surrounding sites with HIV RT, as well as with MMLV RT and bRT-Avd (Fig. S6a). Thus, we limited the abasic site to *TR* 23. HIV RT incorporated adenine almost exclusively (98%) across the abasic site (Fig. 5a), and while deletions still occurred, they occurred much less frequently than with tandem abasic sites (Fig. S6b). MMLV RT has been reported to be intolerant to abasic sites (32), and indeed the incidence of deletion was approximately three-fold higher at abasic *TR* 23 for MMLV RT than for HIV RT (Fig. S6b). Nevertheless, for those cDNAs lacking deletions, MMLV RT incorporated adenine almost exclusively (99%) across the abasic site (Fig. 5a). A significant level of deletion at and around the abasic site was also seen with bRT-Avd, with an incidence that resembled that of MMLV RT (Fig. S6b). However, the near exclusive preference for incorporating adenine seen for HIV and MMLV RTs was not observed for bRT-Avd, and instead at best a strong preference for adenine was evident (65%) (Fig. 5a). Guanine was also incorporated at an appreciable frequency by bRT-Avd across the abasic site (Fig. 5a). These results indicated that bRT-Avd did not follow the A-rule as strictly as HIV RT.

**Figure 5.**
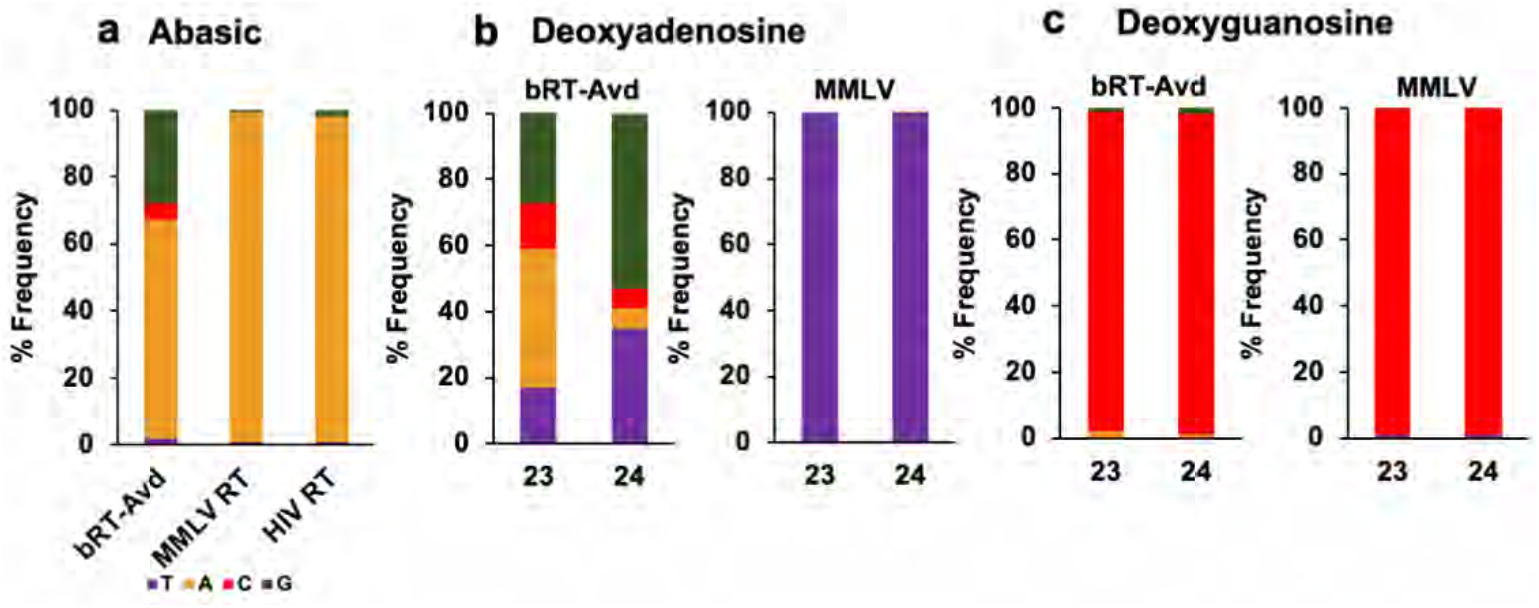
Abasic and Deoxy Template Sites. a. Frequency of deoxynucleotides (mis)incorporated across an abasic site at *TR* 23 for bRT-Avd, MMLV RT, and HIV RT. b. Frequency of deoxynucleotides (mis)incorporated by bRT-Avd (left) and MMLV RT (right) with deoxyadenosines at *TR* 23 and 24. c. Frequency of deoxynucleotides (mis)incorporated by bRT-Avd (left) and MMLV RT (right) with deoxyguanosines at *TR* 23 and 24.

### 2’-OH

We also asked whether adenine-mutagenesis occurred when a deoxynucleotide was reverse transcribed. We therefore instituted deoxyadenosine at *TR* 23 and 24 in the core DGR RNA template. As expected, thymine was incorporated exclusively by MMLV RT across *TR* dA23 and dA24 (Fig. 5b). In contrast, misincorporation increased for bRT-Avd. It went from 40 to 65% at *TR* 24 and from 72 to 83% at *TR* 23 (Fig. 5b). To ascertain whether selectivity to adenine was maintained, *TR* 23 and 24 were instituted with deoxyguanosine. MMLV RT incorporated cytosine (99%) almost exclusively across these sites. Similarly, bRT-Avd incorporated cytosine across deoxyguanosine with nearly the same frequency as across guanosine (97 % vs. 98%) (Fig. 5c). These results indicated that the 2’-OH of a template adenosine promoted correct incorporation and therefore countered adenine-mutagenesis.

### bRT Amino Acids that Modulate Adenine-Mutagenesis

We next sought to identify bRT amino acids that modulated adenine-mutagenesis. We pursued this through structure-guided mutagenesis using a high-confidence model of bRT (100% confidence level for 95% of the amino acid sequence) that was generated using Phyre2 (33) (Supplementary Data). This bRT model was based in part on the structures of group II intron maturases (34,35), including the high fidelity GsI-IIc RT (36). This latter structure also contains a bound RNA template-DNA primer heteroduplex and an incoming dATP. The *in silico* bRT model consisted of all the important functional elements of RTs — the canonical fingers, palm, and thumb domain (Fig. 6a). In addition to this, we relied on the extensive literature on the fidelity of HIV RT, and superposed the structure of HIV RT (37) with the *in silico* model of bRT to guide the choice of substitution sites.

**Figure 6.**
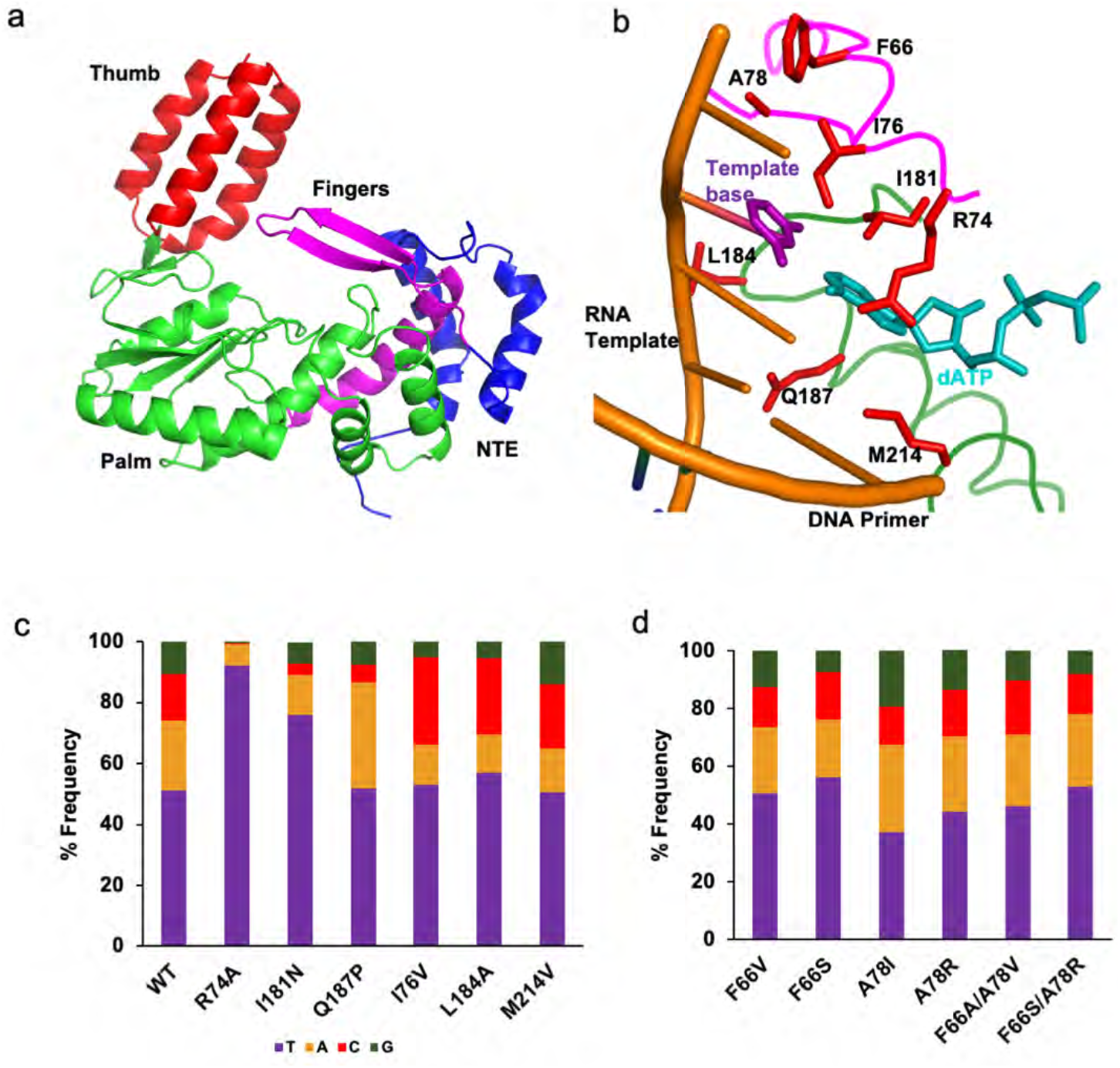
bRT amino acids that modulate selective fidelity. a. *In silico* model of bRT in cartoon representation, showing the N-terminal extension (NTE, blue), Fingers (magenta), Palm (green), and Thumb (red) subdomains. b. *Ιn silico* model of bRT with amino acids subjected to mutagenesis shown as red bonds. The main chain is shown as a coil, colored by subdomains as in panel a. For reference, the RNA template-DNA primer heteroduplex (gold) and incoming dATP (cyan) from the structure of the GsI-IIc group II intron maturase is shown. The template base (thymine) is purple. c. Average frequency of deoxynucleotides (mis)incorporated across the 22 *TR* adenines for wild-type bRT and bRT containing substitutions at amino acids implicated in modulating selective fidelity. d. Average frequency of deoxynucleotides (mis)incorporated across the 22 *TR* adenines for wild-type bRT and bRT containing substitutions at amino acids predicted to be proximal to the templating base.

Based on this superposition, bRT Arg 74 was predicted to form a part of the binding pocket for the incoming dNTP (Fig. 6b). Its putative homolog in HIV RT is Arg 72, which when substituted by Ala decreases misincorporation (38,39). We found a similar effect for bRT(R74A)-Avd. Substitution of bRT Arg 74 with Ala led to a marked decrease in misincorporation frequency across the 22 template adenines in *TR* from 52 to 8%, with some sites reaching as low as 1% misincorporation (Fig. 6c and Fig. S7a). Misincorporation at other template bases (0.4%, 0.4%, and 1.7% for uracil, cytosine, and guanine, respectively) were at the levels within error seen for wild-type bRT (Fig. S7b). Thus, bRT Arg74 is a promoter of adenine-mutagenesis. Single deoxynucleotide primer extension analysis was carried out with bRT R74A to determine the basis for the decreased misincorporation. While there was almost no change in the *K_m_* for the correct incorporation of TTP across a template adenine at *TR* 117, a 40% decrease in *k_cat_* was observed (Table 3). The catalytic efficiency of bRT(R74A)-Avd for proper incorporation was 68% of that of wild-type bRT. Efforts to quantify misincorporation of dATP, the most frequently misincorporated dNTP, by primer extension analysis were unsuccessful, with no misincorporation evident for bRT(R74A)-Avd.

**Table 3.** Steady-state enzymatic parameters for incorporation by mutant bRT-Avd.

| bRT mutant | $K_m$ ($\mu\text{M}$) | $k_{cat}$ ($\text{min}^{-1}$) $\times 10^{-3}$ | $k_{cat}/K_m$ ($\mu\text{M}^{-1} \text{min}^{-1}$) $\times 10^{-3}$ | Efficiency<sup>d</sup> |
| --- | --- | --- | --- | --- |
| R74A, A:T | 7.09 ± 4.68 | 7.03 ± 0.78 | 0.99 | 0.68 |
| I181N, A:T | 218.85 ± 55.94 | 11.13 ± 1.04 | 0.05 | 0.03 |
| I181N, A:dA | 259.10 ± 144.40 | 1.37 ± 0.17 | 0.01 | 0.01 |

<sup>d</sup>Efficiency = $[k_{cat}(\text{mutant})/K_m(\text{mutant})] / [k_{cat}(\text{wt})/K_m(\text{wt})]$

Ile 181 of bRT was also predicted to be proximal to the incoming dNTP, and corresponds to HIV RT Gln 151, whose substitution by Asn decreases misincorporation (40–42). Ile 181 is the first amino acid of the signature DGR RT motif [I/V/L]GxxxSQ (1). In most retroviral RTs, non-LTR retrotransposon RTs, and group II intron maturases, this motif is instead QGxxxSP. We found that bRT I181N decreased the average misincorporation frequency across template adenines from 52 to 24% (Fig. 6c and S7c). The misincorporation frequencies at other template bases were similar within error to those of wild-type bRT-Avd (Fig. S7d, 1.2%, 0.5%, and 3% for uracil, cytosine, and guanine respectively). These results indicated that, like Arg 74, Ile 181 promoted misincorporation across template adenines. Single deoxynucleotide primer extension analysis of bRT(I181N)-Avd for correct TTP incorporation across a template adenine at *TR* 117 showed a large increase in *K_m_* as compared to wild-type bRT-Avd (Table 3, 27-fold). Notably, the *K_m_* for misincorporation of dATP across a template adenine by bRT(I181N)-Avd was similar to that of correct incorporation (Table 3). However, there was a large difference in *k_cat_*. While the *k_cat_*’s for correct incorporation by bRT(I181N)-Avd and wild-type bRT-Avd were similar, the *k_cat_* for misincorporation of dATP by bRT(I181N)-Avd was ~8-fold slower (Table 3). The catalytic efficiency for correct incorporation of TTP by bRT(I181N)-Avd was 3% of wild-type bRT-Avd, and for misincorporation of dATP 13% of wild-type.

We also substituted the last amino acid in the signature DGR RT motif [I/V/L]GxxxSQ, bRT Gln 187, with the proline from the QGxxxSP motif. Proline at this position in HIV RT, P157, contacts a base in the template strand (37). bRT(Q187P)-Avd was unaltered in misincorporation frequency across template adenines compared to wild-type bRT, although this complex showed an increased bias towards misincorporating adenines over other bases (Fig. 6c).

Substitutions were made in three bRT amino acids with putative equivalents in HIV RT that interact with the template or primer strand proximal to the catalytic site (37). These were bRT I176V, L184A, and M214V (Figs. 6b, c). The predicted structural equivalents in HIV RT and the misincorporation-decreasing substitutions are, respectively, Leu 74 substituted with Val (43); Lys 154 substituted with Ala (40); and Met 184 substituted with Val (44,45). While none of these substitutions in bRT changed the misincorporation frequency, they all curiously changed the misincorporation bias towards cytosine and away from adenine (Fig. 6c).

Lastly, we probed two bRT amino acids predicted to be proximal to the template base: Phe 66 and Ala 78 (Fig. 6b). The size and hydrophobicity of these positions were altered through F66V, F66S, A78V, and A78R substitutions, as well as F66A/A78V and F66S/A78R double substitutions. None of these resulted in a significant decrease in misincorporation frequency, and indeed, the misincorporation frequency increased substantially for bRT(A78I)-Avd and somewhat for bRT(A78R)-Avd (Fig. 6d).

## DISCUSSION

DGRs bring about massive protein sequence variation through the unique mechanism of adenine-mutagenesis. This results in variability being restricted to adenine-encoded amino acids, with non-adenine-encoded amino acids remaining conserved. As seen in a number of DGR variable proteins (9,22,46–48), adenine-encoded amino acids are organized by the C-type lectin-fold of the variable protein into a solvent-exposed binding site. Non-adenine-encoded amino acids form the invariable structural scaffolding for the variable binding site. AAY (Y = pyrimidine) codons are especially prevalent in DGR variable proteins, and as previously noted, adenine-mutagenesis of AAY codons captures the gamut of amino acid chemistry but precludes a stop codon (22). Adding a layer of complexity to adenine-mutagenesis is the distribution of misincorporation frequencies documented here. For example, this makes some amino acid positions more variable (e.g., Mtd 357 with A62 in its codon, 76% misincorporation frequency) and some less (e.g., Mtd 369 with A98 in its codon, 30% misincorporation frequency), and thereby shapes the repertoire of ligands functionally bound by DGR variable proteins. These positional effects on misincorporation frequencies are also seen in cDNAs synthesized *in vivo* (13) and *mtd* sequences that have undergone variation in the absence of selection (49). This is most noticeable for *TR* A62, which consistently has a high misincorporation frequency (~70%). No obvious relationship was evident between positional variation in misincorporation frequency and primary sequence. It is possible that the secondary or even tertiary structure of the template plays a role in this.

Extensive work has shown that fidelity in nucleotide polymerases depends both on Watson-Crick hydrogen bonding and shape complementarity between base pairs (50–56). Indeed, shape complementarity appears to be the dominant discriminator (57), as supported by several lines of evidence, including the observation that hydrogen bonding is dispensable for fidelity in DNA polymerase I (58). The general consensus is that pairing between the template and incoming base is sterically evaluated by polymerases (31,52). If the pairing is correct, polymerases undergo an open to closed transition, which places catalytic groups in the right positions for chemistry to proceed. If the pairing is incorrect, the open to closed transition fails to occur, providing time enough for the incorrect dNTP to dissociate before chemistry can occur.

High and low fidelity nucleotide polymerases are alike when it comes to misincorporation but differ crucially with respect to correct incorporation (16). Both types of polymerases display similarly low catalytic efficiencies (*k_cat_/K_m_*) for misincorporation, but high fidelity polymerases have high catalytic efficiencies for correct incorporation while low fidelity polymerases remain at low catalytic efficiencies. High fidelity polymerases have a ~10^5^-fold difference between catalytic efficiencies of correct incorporation versus misincorporation, but low fidelity polymerases have only a ~10^2^-fold difference. While low fidelity polymerases are inefficient enzymes, they have evolved to be inefficient for specific purposes. For example, members of the Y family of DNA polymerases are low fidelity enzymes that are responsible for replicating through DNA lesions (51). Synthesis through a lesion, effectively a non-standard template site, requires low fidelity. Reminiscent of bRT-Avd, the Y family DNA polymerase iota (pol ι) misincorporates dGTP at a frequency of 0.72 across template thymines and has a low catalytic efficiency for incorporating correct base pairs, ranging between 10^−1^ to 10^−4^ μM^−1^min^−1^ (59–61). Notably, the 1-40 x 10^−4^ μM^−1^min^−1^ catalytic efficiency of bRT-Avd for correct incorporation falls within this range (Table 1), and is ~10^4^ lower than the efficiency of high fidelity polymerases (16). DNA pol ι and other Y family DNA polymerases synthesize only short stretches of DNA to repair lesions (62), and indeed DNA pol ι tends to terminate synthesis after incorporating a dGTP across a template thymine (60). This is also reminiscent of bRT-Avd, which synthesizes only short cDNAs (5-35 nt) with non-DGR RNA templates. However, bRT-Avd becomes processive with the DGR RNA as its template and synthesizes extended cDNAs (90- and 120-nt), likely due to the formation of a structured RNP. These observations suggest that the low catalytic efficiency of bRT-Avd is intimately tied to its ability to carry out adenine-mutagenesis, and while the inherent tendency of bRT-Avd is to synthesize short cDNAs, the DGR RNA provides a means to synthesize longer cDNAs. It is possible that a low catalytic efficiency is tolerated by DGRs because the target for variation (e.g., the gene encoding the variable protein) usually exists in single copy number and thus requires only a single cDNA molecule to effect sequence variation. While the efficiency of cDNA synthesis by the *Bordetella* bacteriophage DGR *in vivo* requires further study, it is worth noting that the overall efficiency of sequence variation *in vivo* by this DGR is quite low (10^−6^) (15).

To understand the nucleobase determinants that modulate adenine-mutagenesis, we took advantage of the near isosteric features of infidelity-promoting adenine and fidelity-promoting guanine. Using nucleobase analogs that have adenine- or guanine-like groups, we found that the substituent at the C6 position, but not the N1 or C2 position, had a major effect on misincorporation frequency. An amine at the C6 position, which in adenine acts as a hydrogen bond donor in a Watson-Crick base pair, was functionally equivalent to having no substituent at this position. In contrast, a carbonyl at C6, which in guanine acts as a hydrogen bond acceptor in a Watson-Crick base pair, greatly lowered the misincorporation frequency. Thus, it appears that bRT-Avd is sensitive to the mispositioning of a C6 carbonyl but not a C6 amine.

We also explored the possibility that adenine flips out of the catalytic site, leaving it empty. This has been suggested for DNA pol ι, in which incorporation is more efficient across an abasic rather than a pyrimidine template site (60,61). However, in the case of bRT-Avd an abasic site resulted in a substantial level of deletions, and the misincorporation pattern at the abasic site was not the same as with adenine. While both abasic and adenine sites led predominantly to adenine misincorporation, an abasic site led to preferential misincorporation of guanine over cytosine. In contrast for a template adenine the preference was cytosine over guanine. Thus, the adenine-mutagenesis pattern is not explained by the flipping out of adenine from the catalytic site. In addition, these results indicated that the catalytic site of bRT-Avd is not as predisposed towards an incoming adenine as has been suggested for A-rule polymerases (31).

We sought to identify amino acids in bRT that have a role in modulating adenine-mutagenesis, and relied on a three-dimensional *in silico* model based primarily on group II intron maturases (34–36). We probed eight amino acids predicted to be located at or near the catalytic site, and identified two that modulated adenine-mutagenesis: R74 and I181. Based on an *in silico* model of bRT, Arg 74 is predicted to correspond to Arg 72 of HIV RT. This HIV RT amino acid contacts the base and phosphate of the incoming dNTP (37). Substitution of HIV RT Arg 72 with Ala leads to a significant decrease in misincorporation (about three-fold on average but up to 25-fold at specific sites) with a significant decrease in *k_cat_* (30- to 100-fold) (38,39). Similarly, we observed that substitution of bRT R74 with Ala led to a decrease in misincorporation accompanied by a decrease in *k_cat_*. However, the effects for bRT-Avd were much more modest (1.8-fold decrease in misincorporation and 1.7-fold decrease in *k_cat_*). The second amino acid, Ile 181 of bRT, is predicted to correspond to HIV RT Q151. This amino acid in HIV RT contacts the ribose of the incoming dNTP (37). Substitution of HIV RT Gln 151 with Asn decreases misincorporation by 8- to 27-fold (41), and decreases the affinity for the correct incoming dNTP by 120-fold and for incorrect ones to levels that are not measurable (42). A similar but smaller 1.5-fold decrease in misincorporation across a template was seen for bRT I181N, as well as a 27-fold increase in *K_m_* for correct incorporation of TTP. However, the decrease in misincorporation in bRT(I181N)-Avd was not due to a further increase in *K_m_* for the incorrect deoxynucleotide but instead a marked decrease in *k_cat_*. In HIV RT, Arg 74 and Gln 151 provide contacts that stabilize both correct and incorrect incoming dNTPs. In the absence of the nonspecific contacts provided by Arg 74 and Gln 151, the contributions of correct hydrogen bonding and shape complementarity become more consequential and thereby increase discrimination between pairings, leading to a decrease in misincorporation. Similarly, bRT Arg 74 and Ile 181 are likely to provide nonspecific stabilization of base pairs and thereby promote adenine-mutagenesis.

In summary, our results provide evidence that bRT-Avd is a catalytically inefficient enzyme, a property that is likely intimately tied to its ability to carry out adenine-mutagenesis. We found that the C6, but not the N1 or C2, purine substituent was a key determinant of adenine-mutagenesis, and that Arg 74 and Ile 181 have significant roles in promoting adenine-mutagenesis. Our results provide the first detailed characterization of the nucleobase and protein determinants of adenine-mutagenesis in DGRs.

## DATA AVAILABILITY

The alignment files for NGS data are available at the NIH Sequence Read Archive with accession number PRJNA676165.

## SUPPLEMENTARY DATA

Table S1.

Figures S1-S7.

NGS Script.

bRT Model.

Supplementary Data are available at NAR online.

## ACKNOWLEDGEMENT

We thank Simpson Joseph and Yitzhak Tor for valuable conversations about aspects of this work.

