## Supplemental Table S1 and Figures S1-S7 for "Determinants of Adenine-mutagenesis in Diversity-Generating Retroelements"

Table S1. DNA oligonucleotides

| | |
|---|---|
| P1 | ACCCGAGTTCGACGT GTTGTTCCGCCGAATAGCGCAGCAGCG |
| P2 | ACACTCTTTCCCTACACGACGCTCTTCCGATCTGTCACGCTTTCAGAAGTTAATGGCAAATCTAGACGCTGCTGCGCTATTCGGCG |
| P3 | GACTGGAGTTCAGACGTGTGCTCTTCCGATCTCATTTCCCACAGATTAGCCCCATAATAAAGCTTCAGACGCCGCGCGCCCCGAT |
| P4 | GACTGGAGTTCAGACGTGTGCTCTTCCGATCTATAATAAAGCTTCCCATCACCTTCTTGCATGG |
| P5 | ACCCGGAAGCTTGAAGTCGGCCCCGCCTTTCCA |
| P6 | ATAATAAAGCTTCAGACGCCGCGCGCCCCGAT |
| P7 | GGCAAATCTAGACGCTGCTGCGCTATTCGGCG |
| $P^{117}$ | GAGCCATGCAAGAAGGTGATGGGCA |

Figure S1

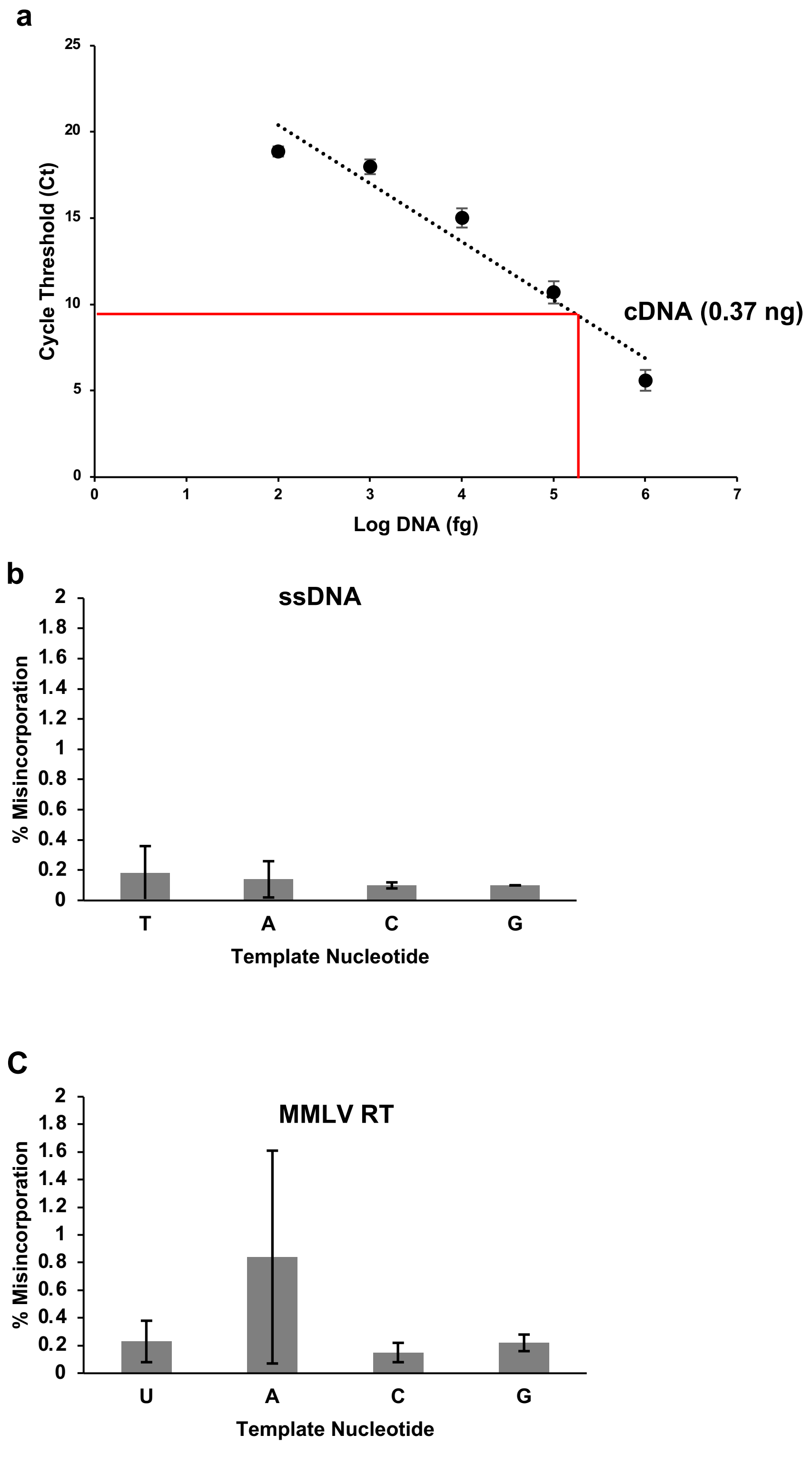


**Figure S1. cDNA quantification**.
**a**. The cycle threshold (Ct) from quantitative PCR reactions plotted against the log of the DNA mass from standards is shown, along with a linear fit. Quantification of the mass of cDNA produced by bRT-Avd is shown by the red lines.
**b**. Misincorporation frequencies from PCR and NGS of chemically synthesized and purified single-stranded DNA at T, A, C, and G template positions. Means and standard deviations are shown.
**c.** Misincorporation frequencies from reverse transcription of DGR core RNA by MMLV RT at U, A, C, and G template positions in *TR*. Means and standard deviations are shown.

Figure S2

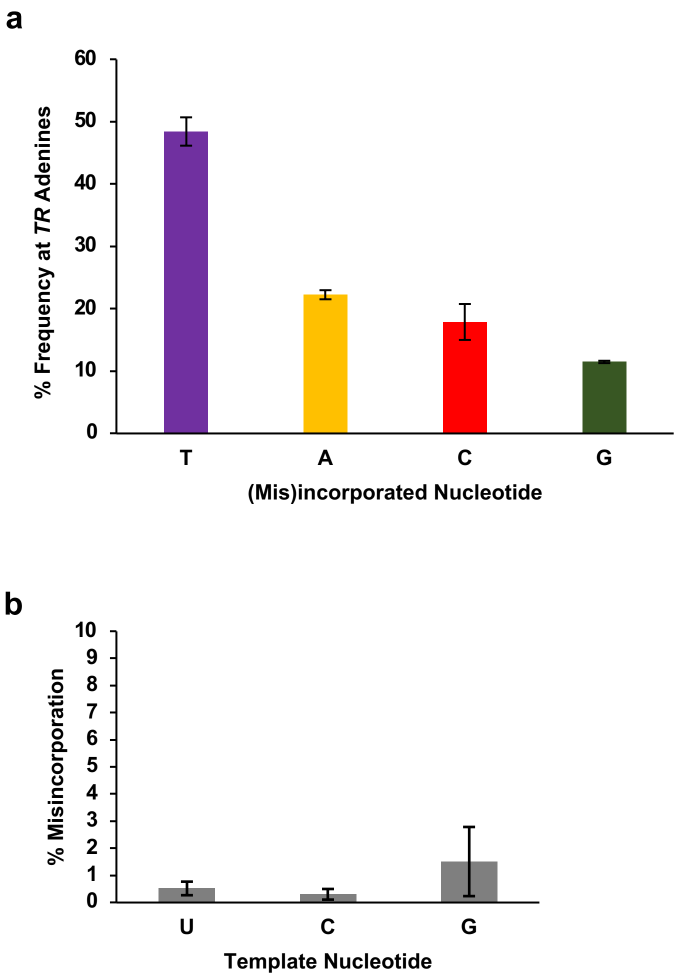


**Figure S2. (Mis)incorporation frequency**.

**a.** Frequency of deoxynucleotide incorporated (thymine) or misincorporated (adenine, cytosine, and guanine) by bRT-Avd across template adenines in *TR*. Means and standard deviations from three independent experiments are shown.

**b.** Frequency of deoxynucleotides misincorporated by bRT-Avd across template uracil, cytosine, and guanine in *TR*. Means and standard deviations from three independent experiments are shown.

Figure S3

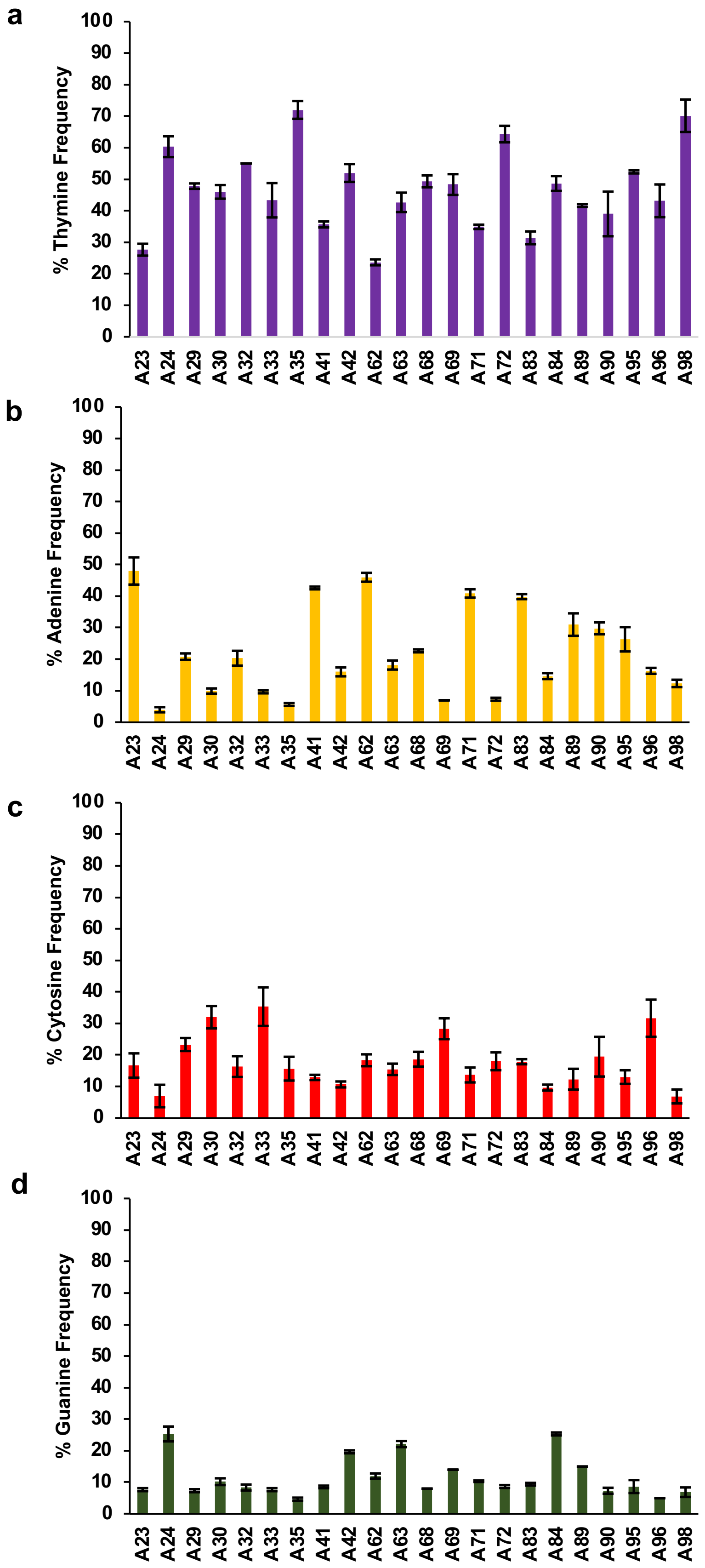
a
% Thymine Frequency
b
% Adenine Frequency
c
% Cytosine Frequency
d
% Guanine Frequency
0
10
20
30
40
50
60
70
80
90
100
A23 A24 A29 A30 A32 A33 A35 A41 A42 A62 A63 A68 A69 A71 A72 A83 A84 A89 A90 A95 A96 A98

**Figure S3. (Mis)incorporation frequency of bRT-Avd**.

Frequency of (**a**) thymine incorporation, and (**b**) adenine, (**c**) cytosine, and (**d**) guanine misincorporation across template adenines in *TR*. Means and standard errors are shown from three independent experiments.

Figure S4

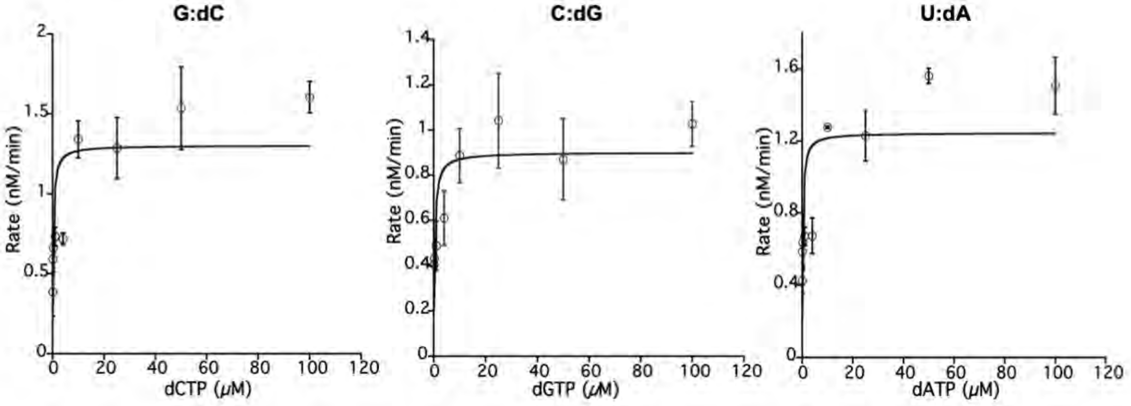


**Figure S4. Kinetics of single deoxynucleotide incorporation**.
Steady-state kinetic characterization of single deoxynucleotide incorporation of dCTP, dGTP, or dATP across guanine, cytosine, and uracil, respectively, at *TR* 117 in the core DGR RNA by bRT-Avd. The Michaelis-Menten fit is shown, and error bars represent standard deviations from three independent measurements.

Figure S5

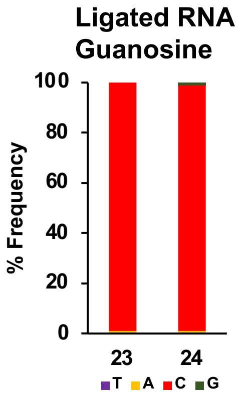


**Figure S5. Ligated RNA with guanosines at *TR* 23 and 24.**

Frequency of deoxynucleotides (mis)incorporated by bRT-Avd at *TR* 23 and 24 using the core DGR RNA template that had been ligated from chemically synthesized and *in vitro* transcribed sections. The chemically synthesized section contained guanosines at *TR* 23 and 24.

Figure S6

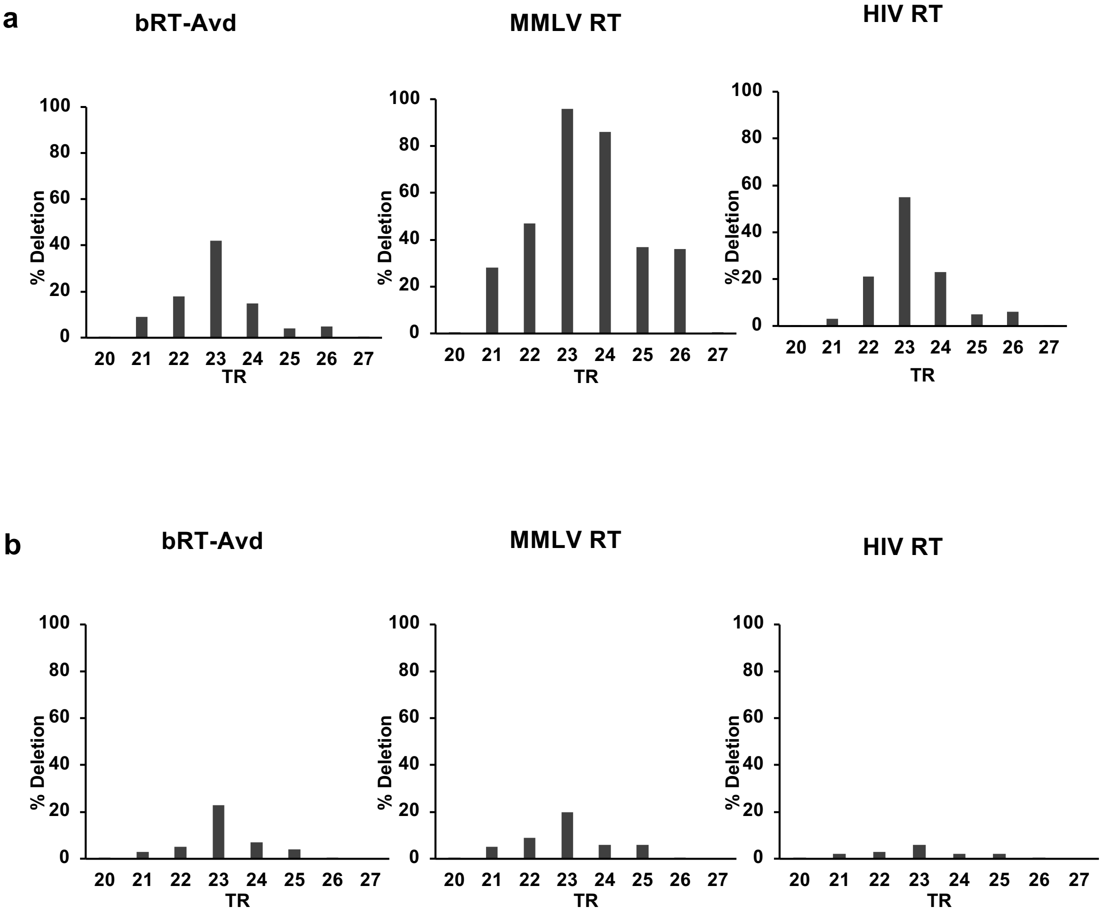


**Figure S6. Abasic sites.**

The frequency of deletions in cDNAs that were reverse transcribed by bRT-Avd, MMLV RT, or HIV RT from the core DGR RNA containing (**a**) tandem abasic sites at *TR* 23 and 24 or (**b**) a single abasic site at *TR* 23.

Figure S7

**a**

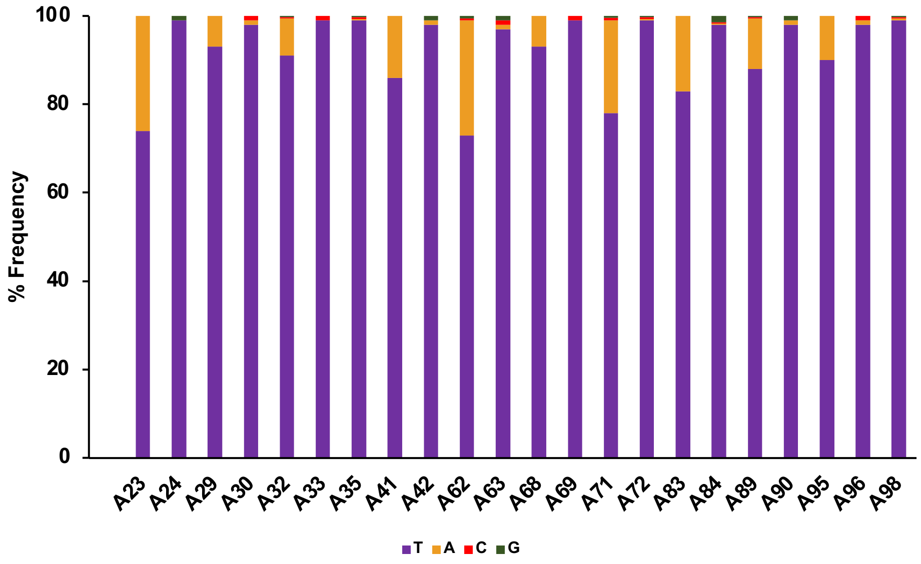


**b**

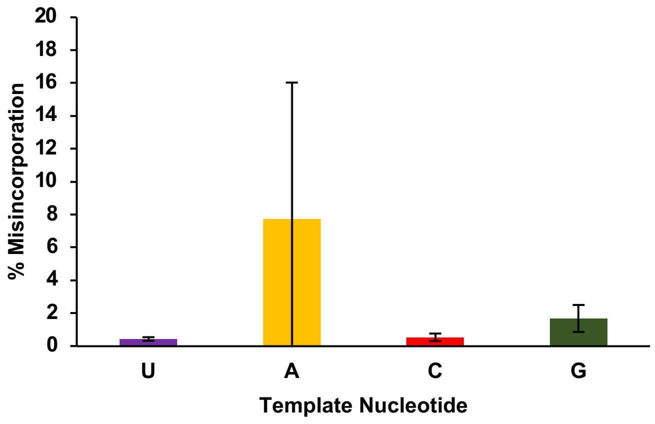


**Figure S7. bRT mutant proteins.**

**a.** Average frequency of deoxynucleotides (mis)incorporated by bRT(R74A)-Avd across template adenines in *TR*.

**b.** Frequency of deoxynucleotides (mis)incorporated by bRT(R74A)-Avd across template uracils, adenines, cytosines, and guanosines in *TR*. Means and standard deviations are shown.

Figure S7 continued

c

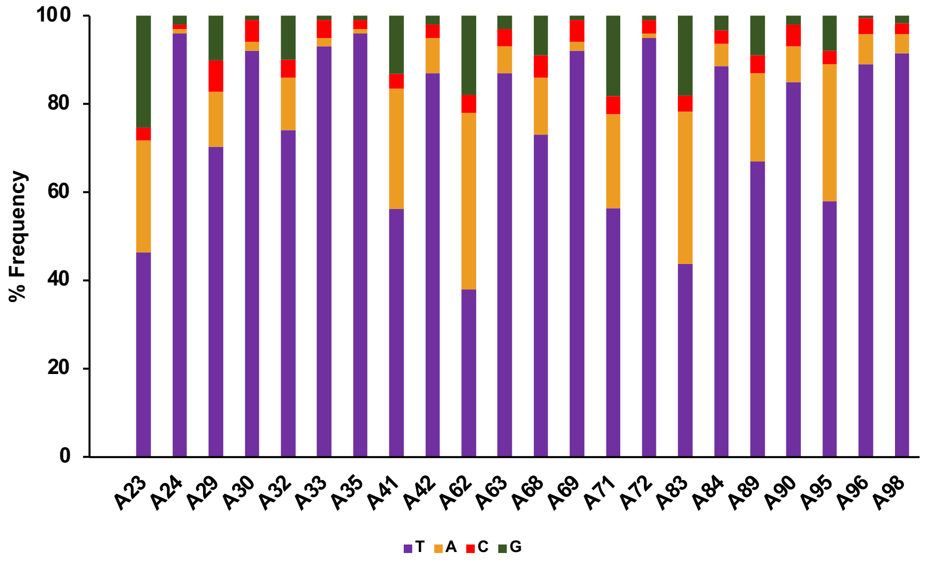


d

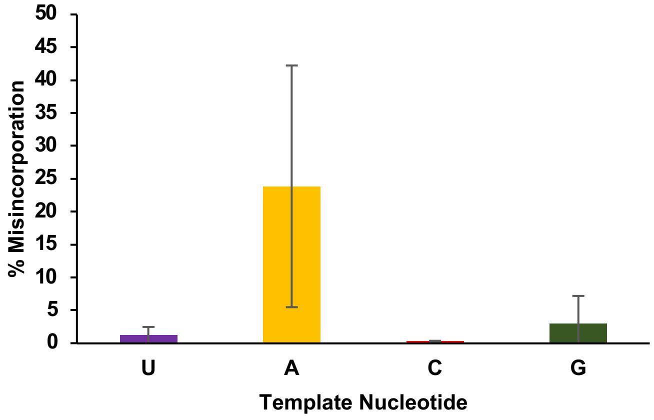


**Figure S7. bRT mutant proteins, continued.**

**c.** Average frequency of deoxynucleotides (mis)incorporated by bRT(I181N)-Avd across template adenines in *TR*.

**d.** Frequency of deoxynucleotides (mis)incorporated by bRT(I181N)-Avd across template uracils, adenines, cytosines, and guanosines in *TR*. Means and standard deviations are shown.

### NGS Scripts

```
# Build a refererence file for bowtie2
# -f flag is for fasta-formatted sequence
bowtie2-build -f ref.fasta bt2ref

# Align the reads to the reference
# -p flag is for how many processes/threads you want to run
# -x is your reference
# -1 is for your paired end fastq #1
# -2 is for paired end fastq #2
# -S is the output file
bowtie2 -p 8 -x bt2ref -1 SH03-G117A-1_S3_L001_R1_001.fastq -2 SH03-
 G117A-1_S3_L001_R2_001.fastq -S alignment.sam

# Converting the .sam file into a sorted and indexed .bam file
samtools view -bS alignment.sam > alignment.bam
samtools sort alignment.bam -o alignment_.sort.bam
samtools index alignment_.sort.bam

# .bam file can be opened in IGV to look at the mutations
```
